# The Hob proteins are novel and conserved lipid binding proteins at ER-PM contact sites

**DOI:** 10.1101/2021.03.02.433623

**Authors:** Sarah D. Neuman, Jeff R. Jorgensen, Amy T. Cavanagh, Jeremy T. Smyth, Jane E. Selegue, Scott D. Emr, Arash Bashirullah

## Abstract

Membrane contact sites are critical junctures for organelle signaling and communication. Endoplasmic reticulum-plasma membrane (ER-PM) contact sites were the first membrane contact sites to be described; however, the protein composition and molecular function of these sites is still emerging. Here, we leverage yeast and *Drosophila* model systems to uncover a novel role for the Hobbit/Hob proteins at ER-PM contact sites. We find that Hobbit localizes to ER-PM contact sites in both yeast cells and the *Drosophila* larval salivary glands, and this localization is mediated by an N-terminal ER membrane anchor and conserved C-terminal sequences. The C-terminus of Hobbit binds to plasma membrane phosphatidylinositols, and the distribution of these lipids is altered in *hobbit* mutant cells. Notably, the Hobbit protein is essential for viability in higher animals, providing one of the first examples of a membrane contact site-localized lipid binding protein that is required for development.

## INTRODUCTION

Eukaryotic cells are defined by the presence of distinct and specialized membrane-bound organelles, and this subcellular compartmentalization enables a wide range of complex cellular functions. In recent years, the importance of organelle membrane contact sites (MCS) for cellular physiology has become increasingly apparent and appreciated; MCS are critical sites for signaling, metabolic channeling, lipid trafficking, organelle fission or fusion, and calcium homeostasis, among other functions (Prinz et al., 2020). MCS are ubiquitous; every type of eukaryotic cell contains them, and every organelle within a cell forms a functional contact site with at least one other organelle (Prinz et al., 2020; Shai et al., 2018; Valm et al., 2017). The endoplasmic reticulum (ER) in particular makes extensive contacts with other organelles, including the plasma membrane (PM), Golgi, mitochondria, peroxisomes, endosomes, lysosomes/vacuoles, and lipid droplets (Wu et al., 2018).

ER-plasma membrane (ER-PM) contact sites were the first MCS to be described. These contact sites were identified in electron microscopy (EM) images of muscle cells in the 1950s (Porter and Palade, 1957). ER-PM contact sites are particularly ubiquitous and prevalent in yeast, with nearly 40% of the PM in contact with the ER (Manford et al., 2012; Pichler et al., 2001; West et al., 2011). Yeast ER-PM contact sites are maintained by six or seven conserved tethering proteins: the tricalbins (Tcb1/2/3), two vesicle-associated membrane protein (VAMP)-associated proteins (VAPs; Scs2 and Scs22), the putative ion channel Ist2, and the more recently described Ice2 (Manford et al., 2012; Quon et al., 2018). Genetic deletion of six tethers (the tricalbins, VAPs, and Ist2; *Δtether* strain) causes dramatic changes in ER morphology, with collapse of cortical ER contacts and redistribution of the ER into the cytosol (Manford et al., 2012). In yeast, ER-PM contact sites play critical roles in lipid trafficking and homeostasis, including phosphatidylinositol trafficking and turnover (Manford et al., 2012; Stefan et al., 2011), sterol trafficking and transfer (Schulz et al., 2009), and maintenance of proper lipid synthesis (Jorgensen et al., 2020; Omnus et al., 2016). However, surprisingly, ER-PM contact sites are not required for yeast viability under normal laboratory growth conditions (Manford et al., 2012).

ER-PM contact sites are also present in metazoans, but the abundance and morphology of these MCS varies widely by cell type (Saheki and De Camilli, 2017). Like yeast, ER-PM contacts in metazoans are maintained by tethering proteins, including the Extended Synaptotagmins (E-Syts; orthologous to yeast tricalbins) (Giordano et al., 2013; Henne et al., 2015), VAPs (Murphy and Levine, 2016), and a number of other proteins that act as dynamic tethers in specific functional contexts or cell types (Eisenberg-Bord et al., 2016; Henne et al., 2015). The functions of ER-PM contact sites in metazoans are diverse but fall into two major categories: calcium homeostasis and non-vesicular lipid transfer.

Store-operated calcium entry (SOCE) is activated to replenish ER calcium stores when they have been depleted (Hogan and Rao, 2015). SOCE is primarily mediated by ER-localized STIM proteins and PM-localized Orai calcium channels, both of which were first discovered in RNAi screens conducted in *Drosophila* S2 cells (Feske et al., 2006; Liou et al., 2005; Roos et al., 2005; Vig et al., 2006; Zhang et al., 2006). SOCE is essential for development, as STIM and Orai knockouts are lethal in both mice and *Drosophila* (Baba et al., 2008; Cuttell et al., 2008; Gwack et al., 2008; Oh-hora et al., 2008; Pathak et al., 2017; Stiber et al., 2008). Several different lipid moieties undergo non-vesicular transfer at ER-PM contact sites, including sterols, glycerophospholipids, and phosphatidylinositols (Henne et al., 2015; Saheki and De Camilli, 2017). However, knockout or mutation of most characterized lipid transfer proteins does not result in animal lethality, suggesting either that lipid transfer is not essential for development or that functionally redundant transfer proteins can compensate for one another.

The Hobbit/Hob proteins are large (>2000 amino acid) proteins that are conserved throughout eukaryotes, but little is known about their molecular function. We identified *hobbit* in a genetic screen for *Drosophila* mutants that arrest development during metamorphosis with a small pupa phenotype, and found that *hobbit* function is required in professional secretory cells, including the insulin producing cells (IPCs) and larval salivary glands, for regulated exocytosis (Neuman and Bashirullah, 2018). Phenotypes have also been observed upon mutation of the plant orthologs of *hobbit*. Mutation of *A. thaliana SABRE* results in short roots with increased diameter and formation of ectopic root hairs; *SABRE* function is thought to be required for proper plant cell expansion and maintenance of planar cell polarity (Aeschbacher et al., 1995; Benfey et al., 1993; Pietra et al., 2015, 2013; Yu et al., 2012). Mutation of *A. thaliana KIP*, a putative paralog of *SABRE*, causes defects in pollen tube growth (Procissi et al., 2003), as does mutation of *Z. mays APT1* (Xu and Dooner, 2006). Finally, mutation of the *SABRE* ortholog in the moss *P. patens* causes stunted growth, defects in polarized growth, and failures in cell division (Cheng and Bezanilla, 2020). Although genetic screens in plants and insects have revealed phenotypic insights, the molecular function of *hobbit* and its orthologs has remained elusive.

Here we use both yeast and *Drosophila* model systems to examine the function of *hobbit*. We find that Hobbit localizes to ER-PM contact sites in both yeast and *Drosophila* salivary gland cells, and this localization is mediated by an N-terminal membrane anchor and conserved C-terminal sequences. Our data also shows that ER-PM localization is required for *hobbit* function. Notably, the C-terminal Apt1 domain of Hobbit binds to phosphatidylinositols, and the distribution of PI(4,5)P_2_ is altered in *hobbit* mutant cells. Together, these results demonstrate that Hobbit is a novel protein that localizes to ER-PM contact sites and suggest that Hobbit may be a lipid transfer protein whose function is required for animal development.

## RESULTS

### The *S. cerevisiae* orthologs of *hobbit* localize to ER-PM contact sites

The *S. cerevisiae* genome contains two predicted orthologs of *hobbit*: *FMP27* and *YPR117W*. To begin to characterize these proteins in yeast, we generated GFP-tagged versions of both proteins under control of their endogenous promoters and examined where they localized within the cell. Both proteins localized in cortical patches or puncta (Fig. 1A, S1A), like other proteins that localize to ER-PM contact sites (Manford et al., 2012). We focused primarily on Fmp27-GFP because Ypr117w-GFP was expressed at barely detectable levels (Fig. S1A). We found that the Fmp27-GFP cortical puncta co-localized with the pan-ER marker RFP-HDEL and with the ER-PM tether Tcb3-mCherry (Fig. 1A-B) (Manford et al., 2012), providing further evidence that Fmp27 localizes to ER-PM contact sites. Some proteins that are enriched at ER-PM contact sites are anchored to the plasma membrane, while others are anchored to the ER (Hogan and Rao, 2015). To distinguish between these possibilities, we examined the localization of Fmp27-GFP in a strain that knocks out six ER-PM tether proteins (*Δtether*) (Manford et al., 2012). Fmp27-GFP was no longer present in cortical puncta in the *Δtether* background and instead localized to perinuclear and internal structures that co-localized with RFP-HDEL (Fig. 1C), indicating that Fmp27 localizes to collapsed ER in the *Δtether* mutant and confirming that Fmp27 is an ER-localized protein. *FMP27* (<u>F</u>ound in <u>M</u>itochondrial <u>P</u>roteome) got its name from mitochondrial proteome analysis (Reinders et al., 2006), raising the possibility that this protein may also be present in mitochondria. We therefore examined the localization of FMP27-GFP with the mitochondrial marker Su9-DsRED but did not observe any co-localization (Fig. S1B), suggesting that Fmp27 may have been a contaminant in the mitochondrial proteome study. Overall, these results demonstrate that Fmp27 is an ER-localized protein enriched at ER-PM contact sites in yeast.

**Figure 1.**
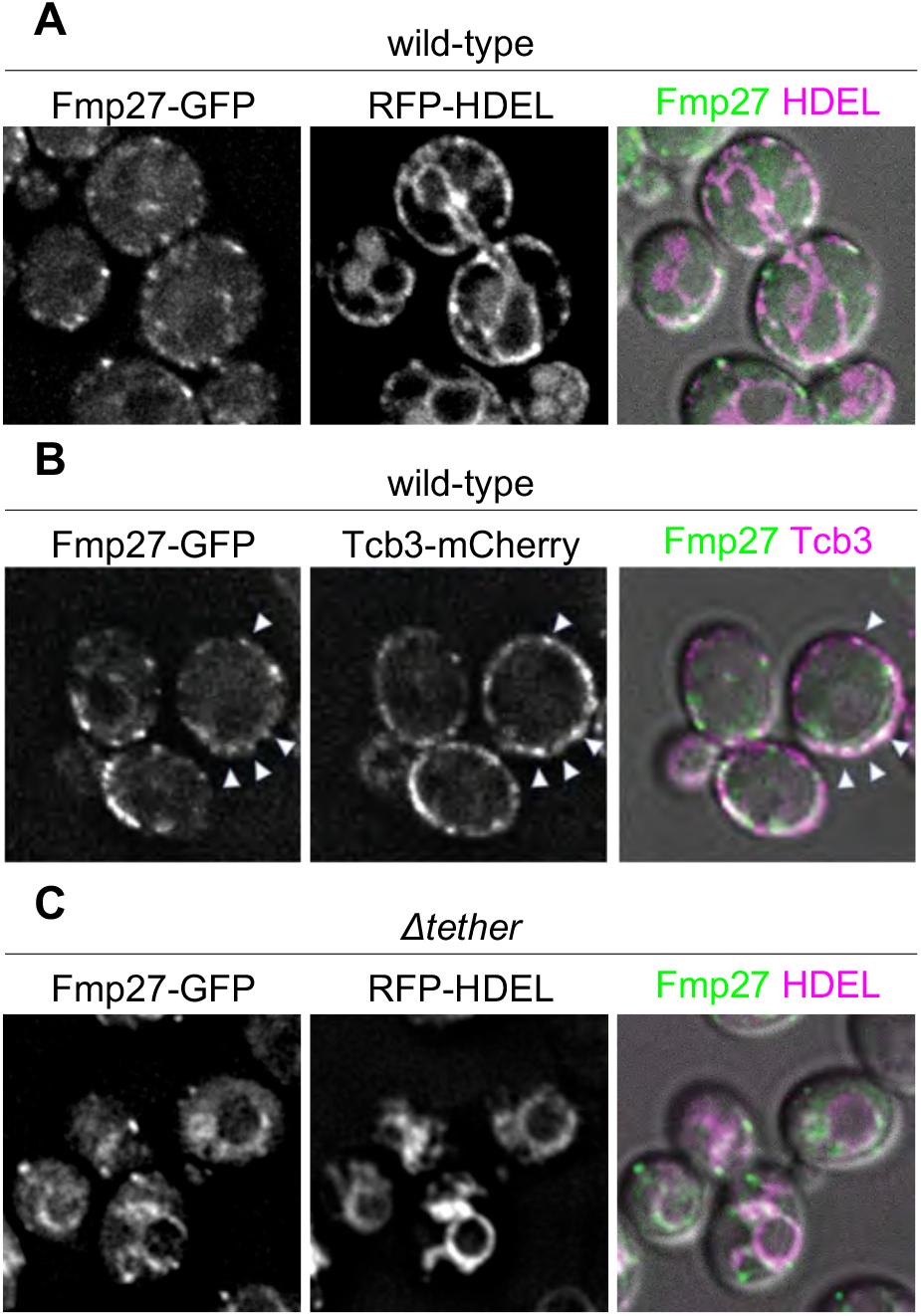
Fmp27 localizes to ER-PM contact sites. **(A)** Live-cell imaging of endogenously tagged Fmp27-GFP (green) shows that Fmp27 is enriched in puncta at the cell cortex that co-localize with RFP-HDEL (magenta), a pan-ER marker. RFP-HDEL is expressed from a plasmid. **(B)** Live-cell imaging of Fmp27-GFP (green) with the ER-PM tether Tcb3-mCherry (magenta) confirms that Fmp27 is enriched at ER-PM contact sites. **(C)** Live-cell imaging of endogenously tagged Fmp27-GFP (green) in the *Δtether* background with RFP-HDEL (magenta) shows that Fmp27 localizes to collapsed ER upon loss of ER-PM tethers. RFP-HDEL is expressed from a plasmid.

### Yeast hobbit localizes to the ER via an N-terminal membrane anchor

We next wanted to determine how Fmp27 localizes to the ER. Bioinformatic transmembrane prediction programs suggest that there may be a transmembrane domain at the N-terminus of the protein. Consistent with this prediction, deletion of the first 192 highly conserved amino acids at the N-terminus of the protein (GFP-Fmp27ΔN192) abolished ER localization and caused the truncated protein to localize to the cytosol (Fig. 2A, S1C), indicating that Fmp27 is anchored to the ER by a transmembrane domain or hairpin at the N-terminus. We next analyzed the membrane topology of Fmp27. We isolated microsomes from cells expressing either Fmp27-GFP or GFP-Fmp27 and performed protease protection assays. In these assays, the luminal domain of Fmp27 should be protected from proteinase K, and GFP, which is a highly stable and tightly folded protein, should run as a free band on a gel if exposed to the cytosol. Note that GFP is resistant to degradation by proteinase K under these experimental conditions. GFP-Fmp27 was barely detectable in its full-length form (Fig. 2B), suggesting that N-terminal GFP tagging interferes with membrane insertion of the protein and may cause it to be degraded. In contrast, Fmp27-GFP was degraded after proteinase K treatment, as evidenced by the appearance of a free GFP band on the gel (Fig. 2B). The ER luminal protein Kar2 (Rose et al., 1989) was used as a control to show that the ER lumen is protected from proteinase K under these conditions (Fig. 2B). These results demonstrate that Fmp27 is anchored to the ER via an N-terminal transmembrane domain or hairpin and the C-terminus of Fmp27 faces the cytosol.

**Figure 2.**
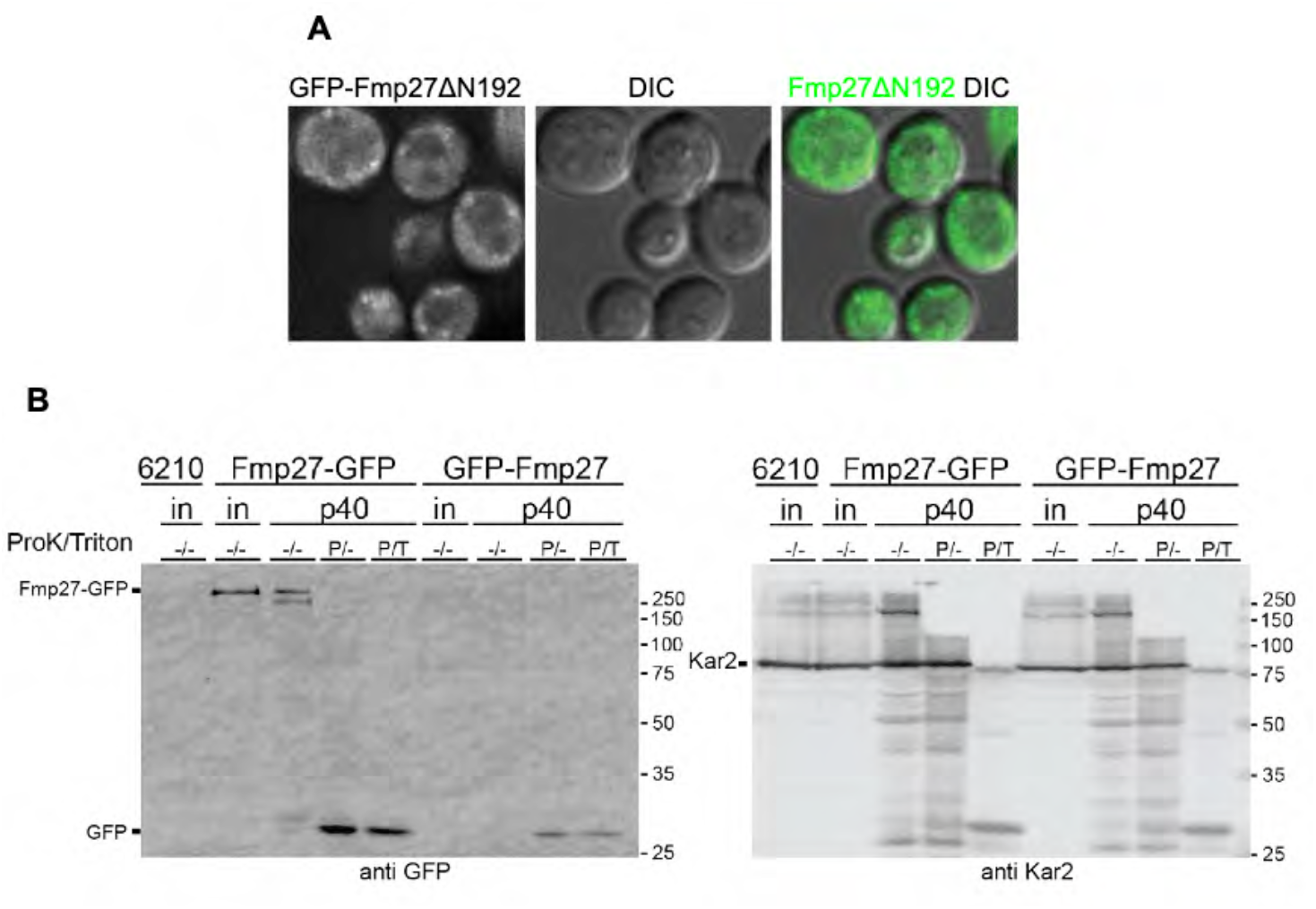
Fmp27 localizes to the ER via an N-terminal membrane anchor with the C-terminus facing the cytosol. **(A)** Live-cell imaging of N-terminally truncated Fmp27 (GFP-Fmp27ΔN192; green) shows that the truncated protein localizes to the cytosol. N-terminal truncation was made at the endogenous FMP27 locus but expressed using the YPQ1 promoter. **(B)** (Left) α-GFP western blot showing Fmp27-GFP and GFP-Fmp27 from cell lysate or the p40 fraction (microsomes). The p40 fraction was either mock-treated, treated with 0.12 µg/mL proteinase K, or 0.12 µg/Ml proteinase K and 1% Triton X-100. The appearance of a free GFP band in the proteinase K-treated Fmp27-GFP p40 fraction shows that the C-terminus of Fmp27 faces the cytosol. (Right) α-Kar2 (ER lumen protein) western blot from the same samples in the left panel. “in” represents input, “P” represents that proteinase K was added, “T” represents that Triton X-100 was added.

### Fmp27 does not function as an ER-PM tether in yeast

Several ER-PM tethering proteins have been identified in yeast, and deletion of six or seven of them is required to significantly reduce the formation of these contact sites (Manford et al., 2012; Quon et al., 2018). However, even upon deletion of seven tethers, cortical ER is still occasionally observed, which could be due to incidental contact or to the function of unidentified tethers (Quon et al., 2018). The topology of Fmp27, with an N-terminal membrane anchor and a large cytosolic domain, coupled with its localization to ER-PM contact sites, is consistent with a potential function as an ER-PM tether. To examine this possibility, we first generated knockouts for both *hobbit* orthologs (*fmp27Δ ypr117wΔ*). *fmp27Δ ypr117wΔ* cells were viable with no obvious growth phenotypes on complete media. Imaging of RFP-HDEL did not reveal any obvious ER morphology defects in *fmp27Δ ypr117wΔ* cells, in contrast to *Δtether* cells, where nearly all cortical ER is lost (Fig. 3A). We also used thin-section transmission electron microscopy (EM) to quantify ER-PM contacts in wild-type, *fmp27Δ ypr117wΔ*, and *Δtether fmp27Δ ypr117wΔ* (8 knockout) mutant cells. No significant differences were observed in the ratio of cortical ER/total PM between wild-type and *fmp27Δ ypr117wΔ* or between *Δtether* and *Δtether fmp27Δ ypr117wΔ* mutant cells (Fig. 3B). We also overexpressed *FMP27* to look for increased ER-PM contacts; however, we observed a slight reduction in the cortical ER/PM ratio in *FMP27*-overexpressing cells (Fig. 3B). Finally, we tested if overexpression of *FMP27* could rescue the loss of cortical ER in *Δtether* cells. We used fluorescence microscopy with RFP-HDEL to examine ER morphology and did not see a rescue of ER-PM contact site formation in *Δtether* cells overexpressing *FMP27* (Fig. 3C). Taken together, these results demonstrate that Fmp27 localizes to ER-PM contact sites but does not appear to function as a tether.

**Figure 3.**
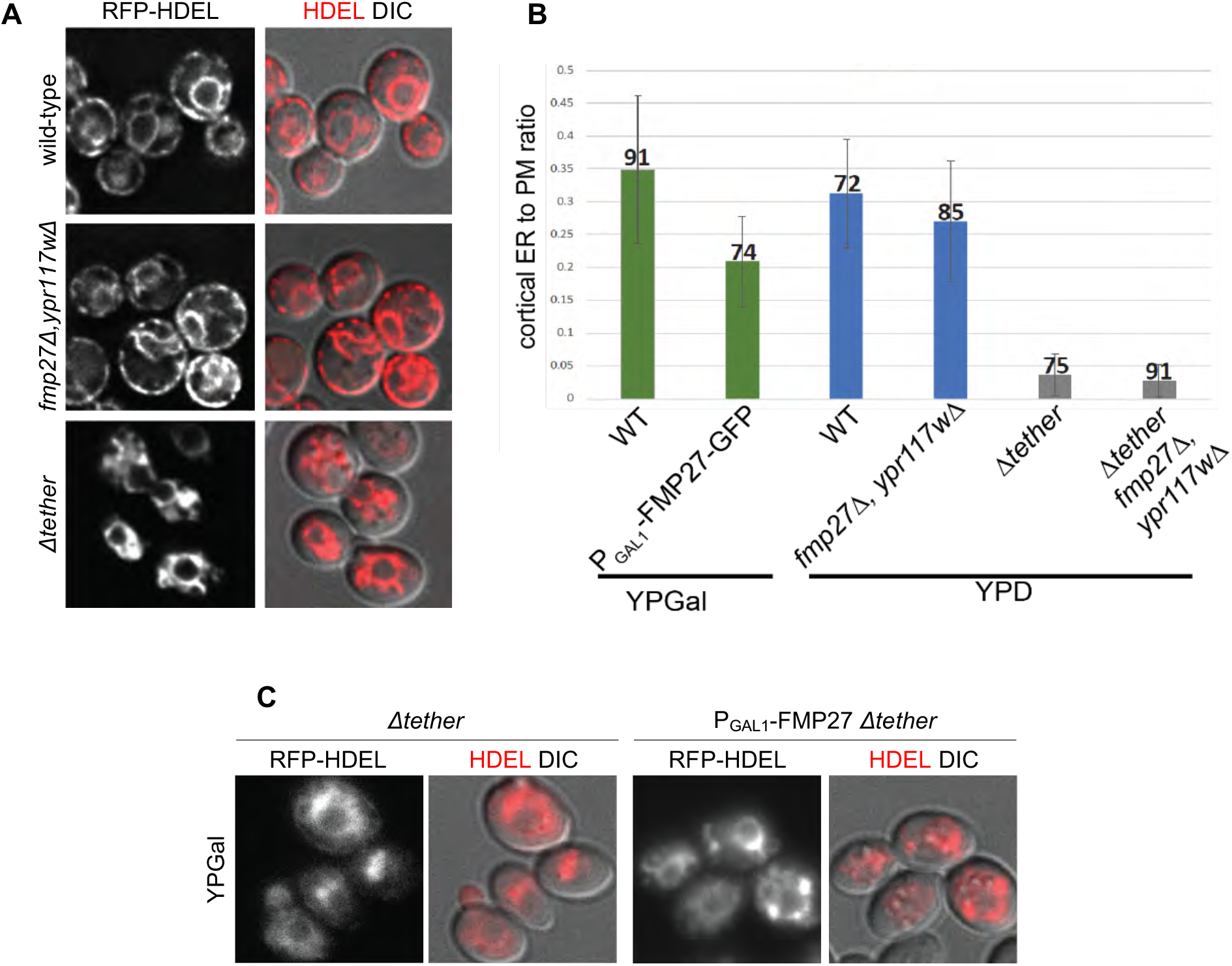
Fmp27 does not function as an ER-PM tether. **(A)** Live-cell imaging of RFP-HDEL (red) in control, *fmp27Δ ypr117wΔ*, and *Δtether* cells shows that ER morphology is unaffected upon loss of both copies of the yeast ortholog of *hobbit*, in contrast to *Δtether* cells, where nearly all cortical ER is lost. **(B)** Quantification of the ratio of cortical ER/total plasma membrane (PM) in control, FMP27-GFP overexpression, *fmp27Δ ypr117wΔ, Δtether*, and *Δtether fmp27Δ ypr117wΔ* shows that overexpression of Fmp27 does not increase the number of cortical ER contacts (green bars), nor does loss of both Fmp27 and Ypr117w decrease the number of cortical ER contacts in a wild-type (blue bars) or *Δtether* background (gray bars). Cells were grown on YPGal media to drive Fmp27 overexpression; otherwise, cells were grown on standard YPD media. Graphs show mean +/-S.D. The number atop each bar represents the number of cells for which the cortical-ER/PM ratio was measured. **(C)** Live-cell imaging of RFP-HDEL (red) in *Δtether* (left) and *Δtether* cells overexpressing Fmp27 (right) shows that overexpression of Fmp27 does not rescue cortical ER contacts in the *Δtether* background. Fmp27 is expressed under its endogenous promoter in the left panels, and under the GAL1 promoter in right panels. Cells were collected from mid-log cultures grown in YPGal media to drive Fmp27 overexpression.

### Membrane topology of *hobbit* is conserved in *Drosophila*

Since we did not observe any obvious phenotypes in yeast that provide clues to the function of *hobbit* at ER-PM contact sites, we turned to *Drosophila*, where mutation of *hobbit* results in a dramatic reduction in animal body size, lethality during metamorphosis, and defects in regulated exocytosis in multiple cell types, including the insulin producing cells and the larval salivary glands (Neuman and Bashirullah, 2018). We first wanted to determine if the ER membrane localization of Hobbit was conserved in flies. Our previous work demonstrated that Hobbit localizes to the ER (Neuman and Bashirullah, 2018). High resolution confocal microscopy enabled us to determine that Hobbit is present on the ER membrane in the larval salivary glands; we observed Hobbit-GFP enriched around the ER luminal marker KDEL-RFP but non-overlapping with cytoplasmic mTagBFP2 (Fig. 4A). Additionally, Kyte-Doolittle hydrophobicity analysis (Kyte and Doolittle, 1982) shows a short, highly hydrophobic region at the N-terminus of the Hobbit protein (Fig. S2A), consistent with a possible transmembrane domain. Like yeast, deletion of a highly conserved N-terminal region resulted in the loss of ER localization, as HobΔN117-GFP co-localized with cytoplasmic mTagBFP2 (Fig. 4B, Fig. S2B). Importantly, ER localization is critical for *hobbit* function, as ubiquitous overexpression of *hobΔN117-GFP* did not rescue the small body size or lethality of *hobbit* mutant animals (Fig. 6C). In contrast, we have previously reported that ubiquitous overexpression of full-length *hobbit-GFP* inserted at the same genomic locus as *hobΔN117-GFP* fully rescues both body size and lethality in *hobbit* mutant animals (Neuman and Bashirullah, 2018). Our next goal was to determine if fly Hobbit had the same membrane topology as yeast Fmp27, and we conducted an imaging-based protease protection assay to analyze the topology. Addition of the selective detergent digitonin, which permeabilizes the plasma membrane but not intracellular membranes (Lorenz et al., 2006), resulted in the release of cytoplasmic mTagBFP2, while Hobbit-GFP and KDEL-RFP were unaffected (Fig. 4C, Movie 1). This further confirms that Hobbit is anchored to the ER membrane. Subsequent addition of proteinase K resulted in the progressive loss of Hobbit-GFP signal, while KDEL-RFP was unaffected (Fig. 4C), confirming that the C-terminus of fly Hobbit faces the cytosol.

**Figure 4.**
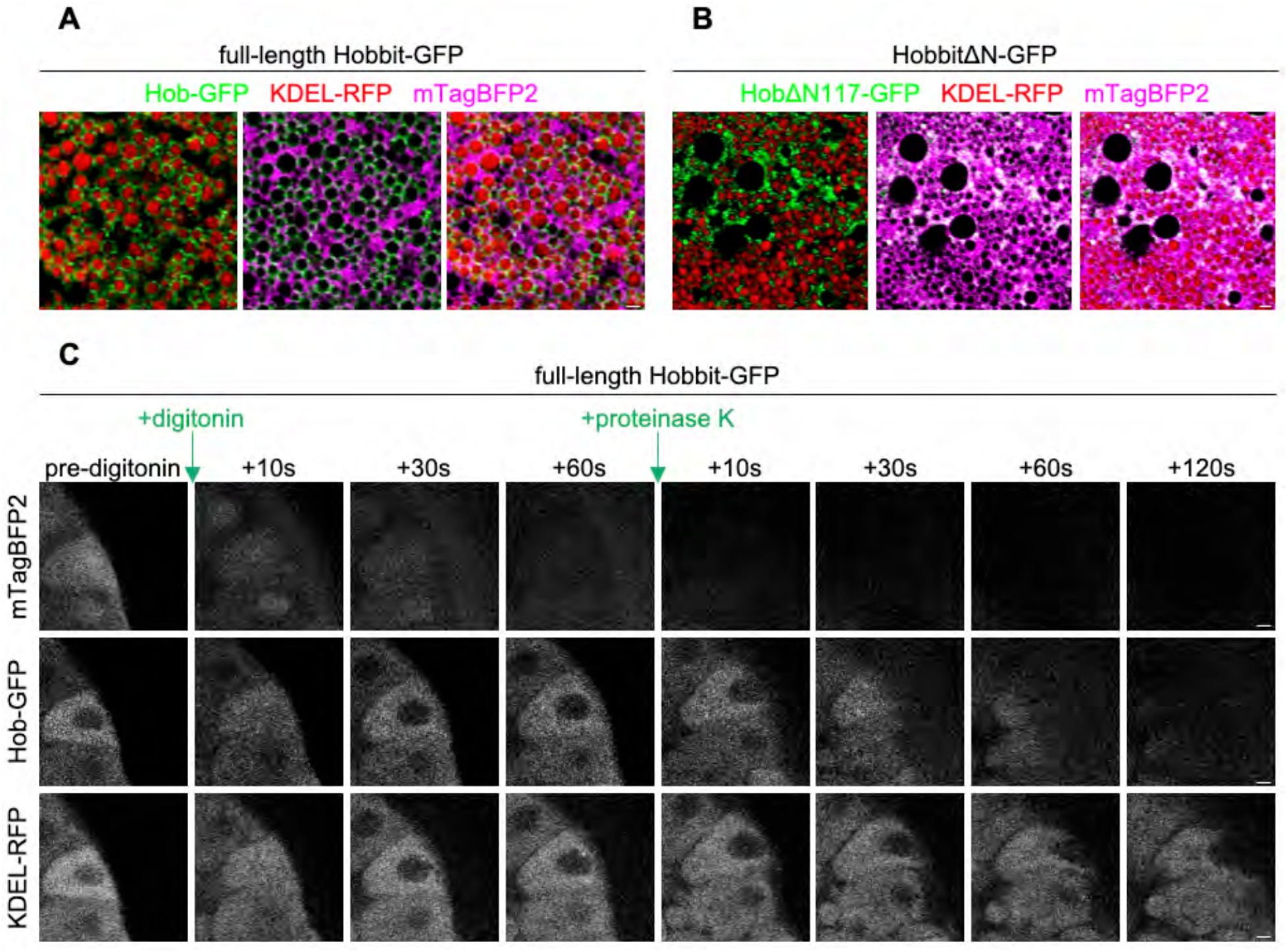
ER membrane localization and topology of Hobbit are conserved in *Drosophila*. **(A)** Live-cell imaging of full-length Hobbit-GFP (green), the ER lumen marker KDEL-RFP (red), and cytosolic mTagBFP2 (magenta) in the *Drosophila* larval salivary glands at the onset of metamorphosis (0 h after puparium formation, PF) shows that Hobbit-GFP localizes to the ER membrane. Full genotype: *UAS-KDEL-RFP/+; Sgs3>hob-GFP/UAS-mTagBFP2*. **(B)** Live-cell imaging of N-terminally truncated HobbitΔN117-GFP (green), the ER lumen marker KDEL-RFP (red), and cytosolic mTagBFP2 (magenta) at 0 h PF shows that HobbitΔN117-GFP localizes to the cytosol. Full genotype: *UAS-KDEL-RFP/ +; Sgs3>hobΔN117-GFP/UAS-mTagBFP2*. Images in (A) and (B) show a single slice from a z-stack comprising three optical sections at a 0.28 µm step size. **(C)** Imaging-based protease protection assay shows that the C-terminus of *Drosophila* Hobbit faces the cytosol. Cytosolic mTagBFP2 (top) rapidly diffuses out of the cells after permeabilization with digitonin, while Hobbit-GFP (middle) and KDEL-RFP (bottom) are unaffected. Hobbit-GFP (tagged at the C-terminus) is degraded after subsequent addition of proteinase K, while KDEL-RFP is unaffected. Note that the cells flatten after addition of proteinase K. Experiment was conducted using 0 h PF glands. Full genotype: *UAS-KDEL-RFP/+; Sgs3>hob-GFP/UAS-mTagBFP2*. Scale bars in (A, B): 1 µm; (C): 10 µm.

### Hobbit localizes to ER-PM contact sites in *Drosophila* cells

Fmp27 is highly enriched at ER-PM contact sites; therefore, we wanted to determine if Hobbit was present at these sites in *Drosophila* cells. The protein composition of ER-PM contact sites is poorly understood in *Drosophila*; therefore, we developed a new marker for these sites based on the structure of the Stim protein. *Drosophila* Stim and its human ortholog STIM1 are well-characterized ER membrane proteins that localize to ER-PM contact sites when ER calcium stores are depleted (Hogan and Rao, 2015). To generate a Stim reagent that constitutively labels ER-PM junctions, we changed D155 and D157 of *Drosophila Stim* (orthologous to D76 and D78 in human STIM1) to A (*Stim*^*DDAA*^*-GFP*). These amino acids are required for calcium binding within the EF hand domain of Stim, and mutation of these sites locks Stim in an extended conformation that enables it to interact with Orai calcium influx channels specifically at ER-PM contact sites (Liou et al., 2005; Lunz et al., 2019; Zhang et al., 2005). Accordingly, we find that Stim^DDAA^-GFP localized in puncta that are present at the plasma membrane in larval salivary gland cells (Fig. 5A), consistent with the expected pattern and localization of ER-PM contact sites. Importantly, full-length Hobbit-mCherry strongly co-localized with Stim^DDAA^-GFP puncta, indicating that fly Hobbit is enriched at ER-PM contact sites (Fig. 5B). Genetic rescue experiments confirmed that the *hobbit-mCherry* transgene produces functional protein (Fig. 6C). Additionally, Stim^DDAA^-GFP appeared to localize normally in *hobbit* mutant cells (Fig. 5A), indicating that ER-PM contact sites still form in the absence of *hobbit*. Taken together, these results demonstrate that Hobbit localization to ER-PM contact sites is conserved between yeast and flies.

**Figure 5:**
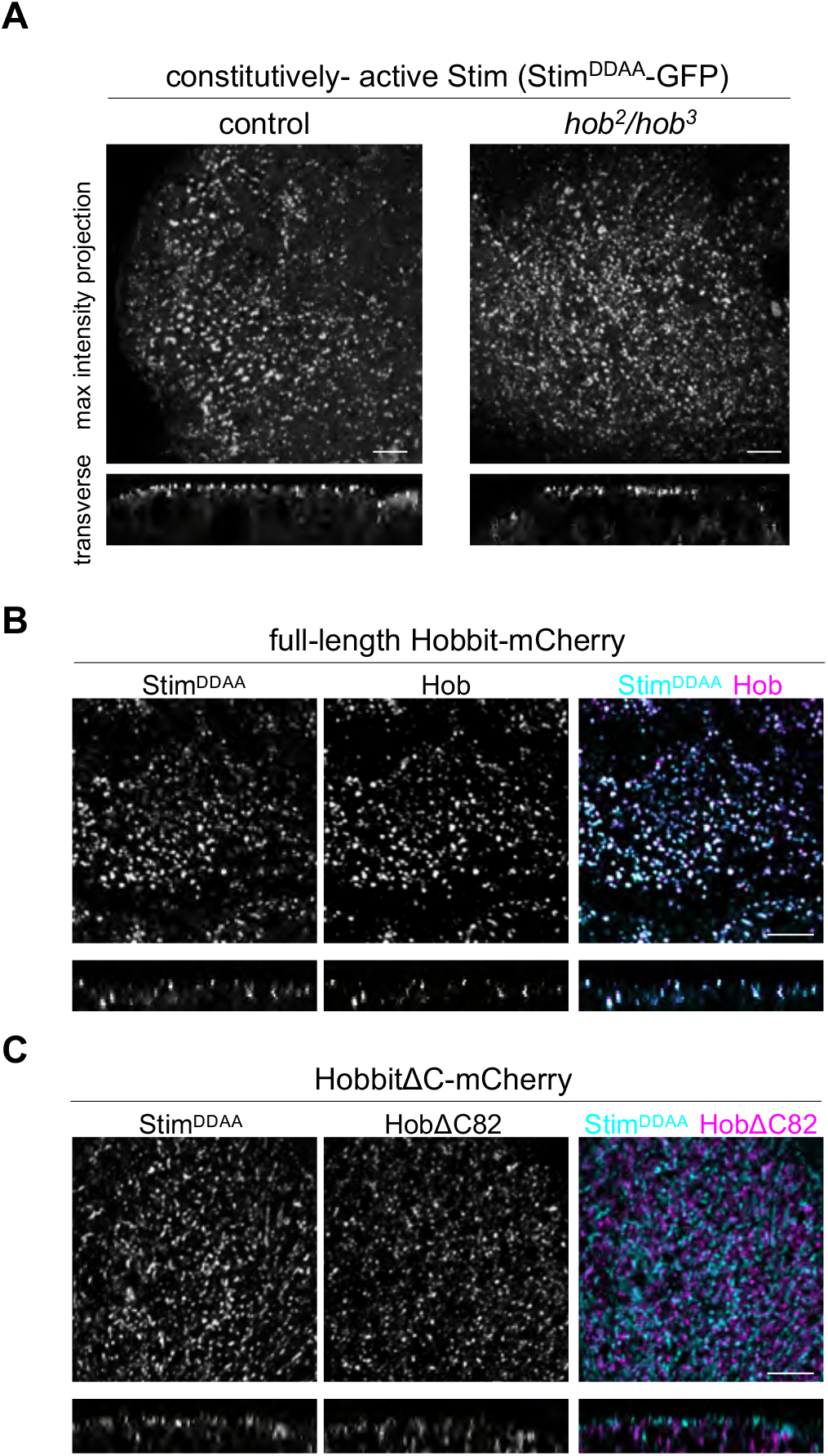
The C-terminus of Hobbit is required for localization to ER-PM contact sites. **(A)** Live-cell imaging of constitutively-active Stim (Stim^DDAA^-GFP) in control and *hobbit* mutant salivary gland cells at the onset of metamorphosis (0 h after puparium formation, PF) shows that this protein localizes to puncta at the plasma membrane. Top images show a maximum intensity projection of 20 optical slices from a z-stack comprising 71 total slices (control) or 57 total slices (*hobbit* mutant) at a 0.36 µm step size. Bottom images show a transverse section from the z-stacks shown above. Full genotypes-control: *UAS-Stim*^*DDAA*^*-GFP/+; Sgs3>/+* and *hobbit* mutant: *UAS-Stim*^*DDAA*^*-GFP/+; hob*^*2*^, *Sgs3>/ hob*^*3*^. **(B)** Live-cell imaging of constitutively-active Stim (Stim^DDAA^-GFP, cyan) and full-length Hobbit (Hobbit-mCherry, magenta) in wandering L3 (wL3) salivary glands shows that Hobbit and constitutively-active Stim co-localize at ER-PM contact sites. Top images show a maximum intensity projection of 10 optical slices from a z-stack comprising 31 total slices at a 0.36 µm step size. Bottom images show a transverse section from the z-stacks shown above. Full genotype: *UAS-Stim*^*DDAA*^*-GFP/+; Sgs3>hob-mCherry/+*. **(C)** Live-cell imaging of constitutively-active Stim (Stim^DDAA^-GFP, cyan) and C-terminally truncated Hobbit (HobbitΔC82-mCherry, magenta) in wandering L3 (wL3) salivary glands shows that HobbitΔC82 no longer localizes to ER-PM contact sites. Top images show a maximum intensity projection of 10 optical slices from a z-stack comprising 31 total slices at a 0.35 µm step size. Bottom images show a transverse section from the z-stacks shown above. Full genotype: *UAS-Stim*^*DDAA*^*-GFP/+; Sgs3>hobΔC82-mCherry/+*. Scale bars: 5 µm.

**Figure 6.**
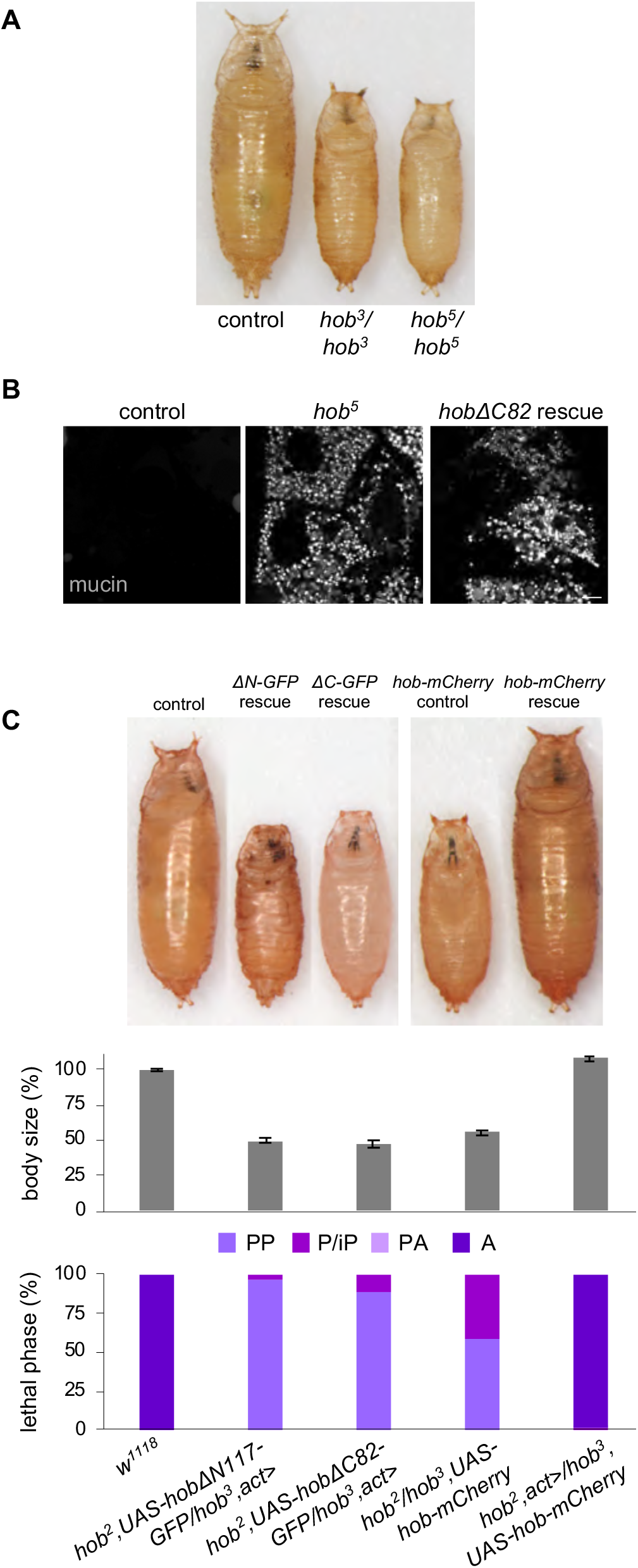
ER-PM localization is required for *hobbit* function. **(A)** Images of control (*w*^*1118*^), homozygous *hob*^*3*^, and homozygous *hob*^*5*^ pupae shows that *hob*^*5*^ mutant animals display the same small body size as *hob*^*3*^ mutant animals. **(B)** Live-cell imaging of mucins in control, homozygous *hob*^*5*^, and *hobΔC82* rescue salivary glands dissected from prepupal animals shows that mucins are secreted in controls but not in *hob*^*5*^ or *hobΔC82* rescue animals. Full genotypes-control: *Sgs3-GFP/+. hob*^*5*^: *Sgs3-GFP/+; hob*^*5*^*/hob*^*5*^. *hobΔC82* rescue: *Sgs3-GFP/+; hob*^*2*^, *UAS-hobΔC82-mCherry; hob*^*3*^, *act>*. **(C)** Body size quantification and lethal phase analysis for control (*w*^*1118*^), *hobΔN117-GFP* rescue (*hob*^*2*^, *UAS-hobΔN117-GFP/hob*^*3*^, *act>*), *hobΔC82-GFP* rescue (*hob*^*2*^, *UAS-hobΔC82-GFP/ hob*^*3*^, *act>*), *hob-mCherry* rescue control (*hob*^*2*^*/hob*^*3*^, *UAS-hob-mCherry*), and *hob-mCherry* rescue (*hob*^*2*^, *act>/hob*^*3*^, *UAS-hob-mCherry*) shows that ubiquitous overexpression of neither N-nor C-terminal truncation of Hobbit rescues the small body size or lethality of *hobbit* mutant animals. In contrast, ubiquitous overexpression of full-length Hobbit-mCherry fully rescues the small body size and lethality of *hobbit* mutant animals. Body size quantified by pupa volume expressed as a percentage relative to control (100%). Data shown as mean +/- S.E.M. *n=*100 animals per genotype. Note that the three pupae on the left were captured in a different image from the two on the right; both were imaged at the same magnification. PP: Prepupa; P/iP: Pupa/incomplete Pupa; PA: Pharate Adult; A: Adult. Scale bar in (B): 10 µm.

### The C-terminus of Hobbit is required for ER-PM localization

We have previously reported five mutant alleles of *hobbit*; all are nonsense mutations scattered throughout the 2300 amino acid protein (Neuman and Bashirullah, 2018) (Fig. S3A). Notably, one of these mutations, *hob*^*5*^, is only 82 amino acids away from the C-terminus of the protein, and this mutant allele is phenotypically indistinguishable from other nonsense mutations located much closer to the N-terminus, with a small body size, lethality during metamorphosis, and cell-autonomous defects in regulated exocytosis in the larval salivary glands (Fig. 6 A-B).

These results suggest that critical functional elements are likely encoded in the final 82 amino acids of the Hobbit protein. C-terminal protein sequences are required for enrichment of other proteins at ER-PM contact sites (Giordano et al., 2013); therefore, we hypothesized that the C-terminal 82 amino acids may be required for Hobbit enrichment at ER-PM contact sites. To test this idea, we generated both GFP- and mCherry-tagged overexpression constructs that delete the final 82 amino acids of the Hobbit protein (*hobΔC82-GFP* and *hobΔC82-mCherry*). These truncated proteins still localized to the ER membrane and topology was unaffected (Fig. S3B-C, Movie 2). However, strikingly, HobΔC82-mCherry did not co-localize with Stim^DDAA^-GFP (Fig. 5C), indicating that ER-PM contact site enrichment was lost upon deletion of the C-terminus of Hobbit. Additionally, ubiquitous overexpression of HobΔC82 did not rescue the small body size or lethality of *hobbit* mutant animals (Fig. 6C), nor did it rescue regulated exocytosis defects in the larval salivary glands (Fig. 6B). We also examined the effect of Fmp27 truncation in yeast cells. Like we observed with fly *hobbit*, deletion of the C-terminus of Fmp27 strongly reduced cortical ER-PM localization (Fig. S4). Overall, these results demonstrate that ER-PM localization is required for *hobbit* function.

### The C-terminal Apt1 domain of Hobbit binds to plasma membrane phosphatidylinositols

Sequence analysis using the protein domain database Pfam (Mistry et al., 2021) predicts that the C-terminus of Hobbit contains an Apt1 domain, a protein domain that is highly conserved but appears to only be present in Hobbit and its orthologs. Primary sequence analysis of yeast Atg2, Vps13, and human VPS13A indicates that these proteins also contain an Apt1 domain (Kaminska et al., 2016; Kolakowski et al., 2020; Rzepnikowska et al., 2017); however, the presence of this domain is not predicted in these proteins by Pfam, perhaps because the primary sequence diverges from the Hobbit Apt1 domain. The Apt1 domains of Atg2, Vps13, and VPS13A bind to phosphatidylinositols (Kaminska et al., 2016; Kolakowski et al., 2020; Rzepnikowska et al., 2017), raising the possibility that Hobbit may also bind to these lipids. To test this hypothesis, we expressed recombinant Hobbit Apt1 domain in *E. coli*, purified it, and tested it for binding on membrane lipid strips. Strikingly, full-length Hobbit Apt1 bound to phosphatidylinositol (PI), PI(4)P, PI(4,5)P_2_, and PI(3,4,5)P_3_, while Apt1 lacking the C-terminal 82 amino acids (Apt1ΔC82) did not bind to any of these lipids (Fig. 7A). These lipid moieties are known to be enriched at the plasma membrane (Balla, 2013), suggesting that the ER-PM localization of Hobbit may be mediated by binding to plasma membrane phosphatidylinositols. We also examined the subcellular distribution of one of these lipids, PI(4,5)P2, in control and *hobbit* mutant cells using the fluorescently-tagged binding reporter PLCδPH-GFP (Verstreken et al., 2009). As expected, PLCδPH-GFP was strongly enriched at the plasma membrane in both control and *hobbit* mutant cells; however, PLCδPH-GFP also accumulated in large intracellular compartments in *hobbit* mutant cells (Fig. 7B), suggesting that *hobbit* plays a functional role in regulating the subcellular distribution of PI(4,5)P_2_.

**Figure 7:**
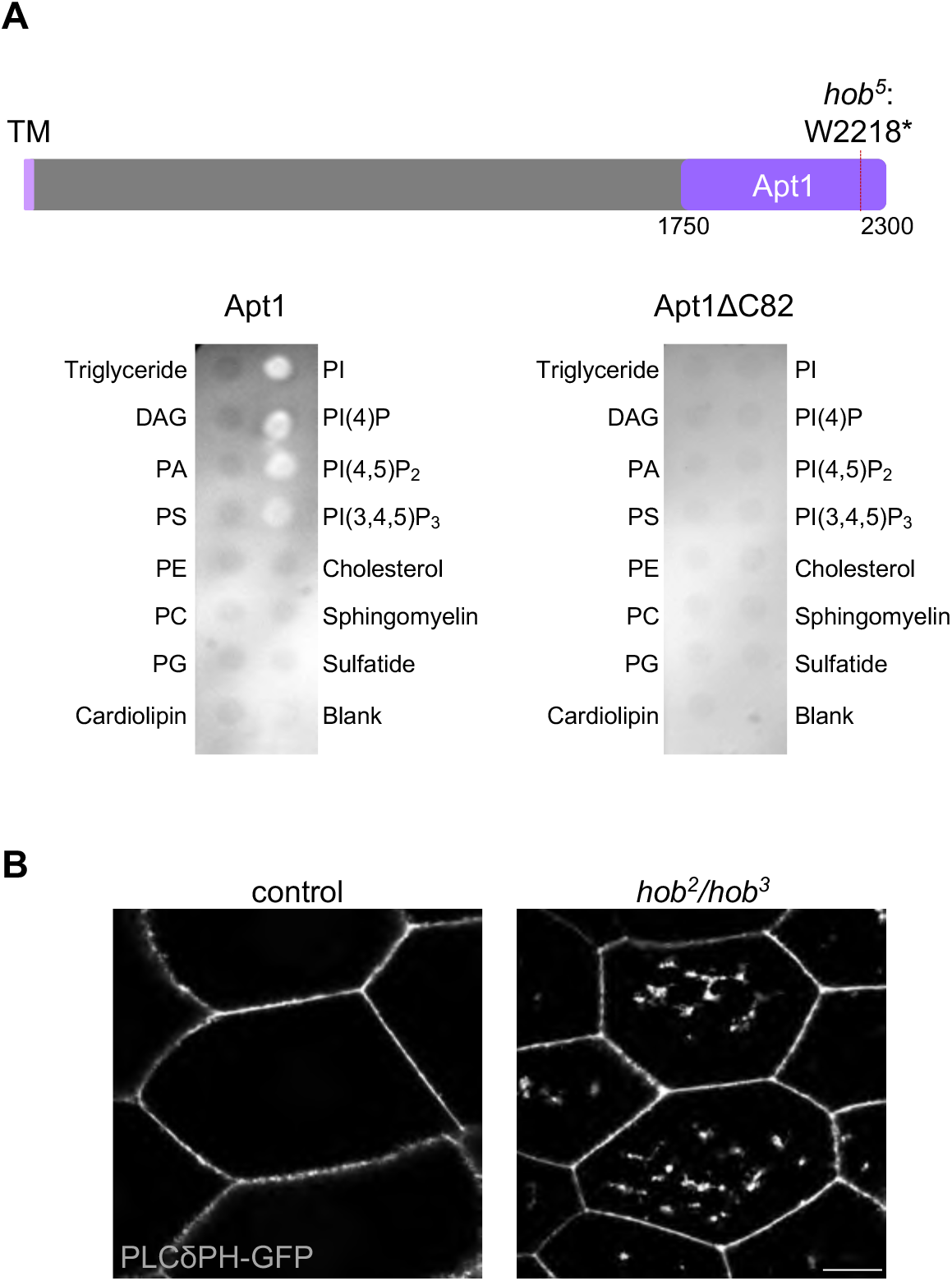
Hobbit binds to phosphatidylinositols and affects PI(4,5)P_2_ distribution. **(A)** Top: schematic depicting the Apt1 domain of Hobbit and the location of the *hob*^*5*^ nonsense mutation. Note that Pfam annotates the Apt1 domain as amino acids 1774-2239, but we included extra upstream and downstream sequence to ensure proper folding of the domain. Bottom: Lipid blot binding analysis shows that full-length Apt1 binds to phosphatidylinositol (PI), PI(4)P, PI(4,5)P_2_, and PI(3,4,5)P_3_, while Apt1ΔC82 does not bind to any of these lipids. Both blots were imaged at the same time with identical acquisition parameters. DAG: diacylglycerol; PA: phosphatidic acid; PS: phosphatidylserine; PE: phosphatidylethanolamine; PC: phosphatidylcholine; PG: phosphatidylglycerol. **(B)** Live-cell imaging of the PI(4,5)P_2_ marker PLCδPH-GFP in control and *hobbit* mutant wandering L3 (wL3) salivary gland cells shows that this lipid accumulates in large intracellular compartments in *hobbit* mutant cells. Full genotypes-control: *Sgs3>/UAS-PLCδPH-GFP* and *hob* mutant: *hob*^*2*^, *Sgs3>/hob*^*3*^, *UAS-PLCδPH-GFP*. Scale bar 20 µm.

## DISCUSSION

ER-PM contact sites are critical junctures for calcium and lipid homeostasis in all eukaryotic cells. Here we have identified Hobbit as a novel protein that is enriched at ER-PM contact sites in both yeast and *Drosophila*. The topology of the Hobbit protein and the requirement for specific C-terminal sequences in ER-PM localization are also evolutionarily conserved. Our data demonstrates that the C-terminal Apt1 domain of Hobbit binds directly to phosphatidylinositols, and *hobbit* appears to play a role in regulating the subcellular distribution of these lipids. Loss of *hobbit* function results in lethality in *Drosophila*; therefore, *hobbit* represents a novel example of a lipid trafficking regulator that is essential for animal development.

The Pfam database (Mistry et al., 2021) annotates the Apt1 domain as a Golgi body localization domain. This annotation comes from analysis of the subcellular localization of a short C-terminal fragment of *Z. mays APT1*, which co-localized with a Golgi marker *in vivo* (Xu and Dooner, 2006). However, our data, coupled with analysis of yeast Atg2 and Vps13 and human VPS13A (Kaminska et al., 2016; Kolakowski et al., 2020; Rzepnikowska et al., 2017), suggests that the Apt1 domain represents a novel phosphatidylinositol binding domain. PI(4)P, one of the lipids that Hobbit Apt1 bound to in our study, is most highly enriched in the plasma membrane and the Golgi (Vermeer et al., 2009), suggesting that the *Z. mays* protein fragment may have been enriched at the Golgi due to PI(4)P binding. Interestingly, the specificity for phosphatidylinositols appears to change among divergent Apt1 domains. Fly Hobbit Apt1 bound to PI, PI(4)P, PI(4,5)P_2_, and PI(3,4,5)P_3_, while yeast Vps13 bound to PI(3)P and human VPS13A bound to PI(3)P and PI(5)P (Kolakowski et al., 2020; Rzepnikowska et al., 2017). Thus, Apt1 domains may represent a novel family of phosphatidylinositol binding domains with varying specificity.

Given that Hobbit binds to phosphatidylinositols and that PI(4,5)P_2_ distribution appears to be altered in *hobbit* mutant cells, our data suggests that *hobbit* may be a novel lipid transfer protein at ER-PM contact sites. Hobbit also exhibits homology to Atg2 and Vps13 in the Apt1 domain; both of these proteins mediate lipid transfer at membrane contact sites (Kumar et al., 2018; Li et al., 2020; Maeda et al., 2019; Osawa et al., 2019; Valverde et al., 2019). Several phosphatidylinositol transfer proteins (PITPs) that function at ER-PM contact sites have previously been characterized. When phospholipase C (PLC) is activated, it hydrolyzes PI(4,5)P2 to generate inositol 1,4,5,-trisphosphate (IP_3_) and diacylglycerol (DAG), which is subsequently converted into phosphatidic acid (PA) (Michell, 1975). PITPs are localized in the cytoplasm but contain domains that allow them to interact with VAPs on the ER membrane and lipids at the PM; upon hydrolysis of PI(4,5)P_2_, PITPs transfer PI from the ER membrane to the PM and PA from the PM to the ER membrane, thereby replenishing the PM store of PI for new synthesis of PI(4,5)P_2_ and allowing recycling of PA in the ER (Cockcroft et al., 2016; Kim et al., 2015; Milligan et al., 1997; Saheki and De Camilli, 2017; Yadav et al., 2015). Hobbit is different from known PITPs in that it is anchored in the ER membrane and binds to both ER-enriched PI as well as to PM-enriched PI(4)P, PI(4,5)P_2_, and PI(3,4,5)P_3_. Future work will be required to directly test if Hobbit functions as a lipid transfer protein *in vitro* and, if so, which lipids are transferred by Hobbit. If *hobbit* does function as a lipid transfer protein, it would be the first such example of a lipid transfer protein that is essential for animal development.

The most conspicuous phenotype in *Drosophila hobbit* mutant animals is a dramatic reduction in body size caused by failure to secrete insulin from the IPCs; *hobbit* mutant animals also exhibit defects in regulated exocytosis of mucin-like ‘glue’ proteins from the larval salivary glands (Neuman and Bashirullah, 2018). Interestingly, mutation of the *Z. mays* and *A. thaliana* orthologs of *hobbit* causes defects in root hair tip growth and pollen tube growth, and both of these processes are highly dependent upon secretion (Guan et al., 2013; Procissi et al., 2003; Xu and Dooner, 2006). Thus, there appears to be an evolutionarily shared requirement for *hobbit* function in cells with high secretory loads. However, our data presented here suggests that the molecular function of *hobbit* lies in the regulation of phosphatidylinositol homeostasis, indicating that the effects on secretion may be indirect. Phosphatidylinositols play essential roles in regulating intracellular trafficking, both in defining membrane identity and in ensuring the recruitment of the proper subset of proteins to the appropriate organelle membrane (Balla, 2013); therefore, disruption of phosphatidylinositol subcellular distribution would be expected to interfere with membrane trafficking. We do not observe widespread membrane trafficking defects in *hobbit* mutant cells; instead, regulated exocytosis appears to be specifically affected. The impact of phosphatidylinositol dynamics on regulated exocytosis in *hobbit* mutant cells remains an important open question.

## MATERIALS AND METHODS

### Yeast media and genetics

Standard recipes were used for YPD and synthetic drop out media (Sherman, 2002). Drop out media was used for selection and retention of plasmids. All imaging experiments were done using synthetic complete or drop out media. Gene deletions and GFP-fusions were made following standard genetic techniques (Longtine et al., 1998).

### Yeast imaging

1 mL of mid-log cultures grown in synthetic media were spun down, and all but ∼50 µL of media were aspirated. The pellet was then re-suspended in the remaining media, resulting in a concentrated suspension of cells. Cells were then imaged on a DeltaVision RT system, using a DV Elite CMOS camera and a 100x objective. Image deconvolution was performed using DeltaVision software (softWoRx 6.5.2).

### Yeast protease protection assay

40 OD of cells were collected from mid-log cultures grown in YPD. Cell pellets were re-suspended in pre-spheroplast media (100 mM Tris pH 8, 10 mM DTT) and incubated for 10 min at room temperature. Cells were spun down and re-suspended in 4 mL spheroplast buffer (0.1 M phosphocitrate, 1.0 M sorbitol, 0.5 mg/mL zymolyase) and incubated at 30°C for 30 min. Cells were then spun down and washed with 4 mL 0.1 M phosphocitrate, 1 M sorbitol. To lyse cells, cells were spun down and re-suspended in 4.3 mL lysis buffer (50 mM Tris pH 7.5, 0.6 M sorbitol), then the suspension was dounced 15 times. The lysis product was spun at 500x*g* for 5 min to remove intact cells, and 200 uL of the supernatant was taken as the input sample. The remaining supernatant was centrifuged at 40,000x*g* for 10 min. The supernatant was discarded, and the pellet was resuspended in 100 µL lysis buffer. The resuspended pellet was then split into 3 tubes, 33 µL each. Tube one was used as a mock experiment. The second tube was the protease protection experiment, and contained 0.12 µg/mL proteinase K. The third tube was the protease positive control experiment and contained 0.12 µg/mL proteinase K and 1% Triton X-100 to disrupt membranes. The tubes were incubated for 25 min at room temperature. To destroy proteinase K function, 3 µL 0.2 M PMSF was added, followed by 33 µL 2x sample buffer (150 mM Tris HCl, pH 6.8, 7 M urea, 10% SDS, 24% glycerol, bromophenol blue). The input fraction was precipitated in 10% TCA and re-suspended in 100 µL 2x sample buffer. Samples were heated at 42°C for 7 min then loaded on an 11% SDS page gel for western blotting using standard procedures. Mouse anti-GFP (B2) (Santa Cruz sc-9996) was diluted to 1:500 and rabbit anti-Kar2 (a gift from R. Schekman, University of California at Berkeley) was diluted to 1:20,000.

### Electron microscopy and analysis of ER-PM contact sites in yeast

Yeast cells were grown overnight in YPD or YPGal and harvested during mid-log growth phase. Cell fixing and staining procedure were performed as previously described (Jorgensen et al., 2020). For quantification of ER-PM contacts, the researcher was blinded to the strain and growth conditions of cells in the EM images. ImageJ was used to measure the length of the PM from each cell by tracing the PM using the freehand line tool, then measuring the length of each ER-PM contact site using the same tool. The sum of the ER-PM contact measurements was then divided by the total length of the PM of each image. These values were used to calculate the average cER/PM ratio for each strain.

### Sequence alignments and conservation analysis

To identify highly conserved protein sequences in Fmp27, the *S. cerevisiae* Fmp27 protein sequence was aligned with the orthologous protein sequence from *K. naganishii, T. blattae, K. capsulate, C. albicans, C. beticola, N. crassa, U. maydis, A. flavus*, and *S. pombe*. The multiple sequence alignment was performed using MAFFT version 7 (Katoh et al., 2018; Kuraku et al., 2013). For the *Drosophila* protein, an alignment was performed between *D. melanogaster hobbit* and its orthologs in *D. willistoni, D. ananassae, D. erecta, D. simulans, D. sechellia, D. grimshawi, D. virilis, D. mojavensis*, and *D. busckii* using ClustalOmega (Sievers et al., 2011). For both yeast and *Drosophila*, the alignments were visualized in Jalview (Waterhouse et al., 2009) and conservation scores were extracted at each position to generate the identity plots.

### Drosophila stocks, husbandry, and body size/lethal phase analysis

The following fly stocks were obtained from the Bloomington *Drosophila* Stock Center: *UAS-KDEL-RFP, UAS-mTagBFP2, Sgs3-GAL4, Sgs3-GFP, act-GAL4, UAS-PLCδPH-GFP. UAS-hobbit- GFP* and the *hobbit* mutant alleles were previously described (Neuman and Bashirullah, 2018). We have used the “>“ symbol in genotypes as shorthand for “GAL4.” All experimental crosses were grown on standard cornmeal-molasses media in uncrowded bottles or vials kept in an incubator set to 25°C. Body size quantification was performed as previously described (Neuman and Bashirullah, 2018). Pupa images were captured on an Olympus SXZ16 stereomicroscope coupled to an Olympus DP72 digital camera with DP2-BSW software. For lethal phase analysis, pupae were allowed to age on grape agar plates at 25°C for one week; then, Bainbridge and Bownes staging criteria (Bainbridge and Bownes, 1981) were used to determine lethal phase.

### Drosophila transgene generation

To generate *UAS-hobbitΔC82-GFP*, the *pENTR 1A-hobbit* plasmid (Neuman and Bashirullah, 2018) was digested with *Spe*I and *Not*I. The following primers were annealed, phorphorylated, and ligated into *pENTR 1A-hobbit*: 5’-CTA GTC ATA CCT ACG CTG GAG TAT CAC AAT GTG ACG AAG C-3’ and 5’-GGC CGC TTC GTC ACA TTG TGA TAC TCC AGC GTA GGT ATG A-3’. The resulting plasmid (*pENTR 1A-hobbitΔC82*) was sequence-verified and recombined into the Gateway cloning destination vector *pBID-UASC-G-GFP* (Neuman and Bashirullah, 2018; Wang et al., 2012) using LR Clonase (Invitrogen). Successful recombination was confirmed by sequencing. To generate *UAS-hobbitΔC82-mCherry, mCherry* was first amplified from *pLV-mCherry* (Addgene) using the following primers: 5’-ACA GGT ACC AGT GAG CAA GGG CGA GGA GGA T-3’ and 5’-ACA TCT AGA TAC TTG TAC AGC TCG TCC ATG-3’. The resulting PCR product was digested with *Kpn*I and *Xba*I and ligated into *pBID-UASC-G-GFP* (thereby replacing GFP with mCherry). The *pENTR 1A-hobbitΔC82* entry clone was recombined *into pBID-UASC-G-mCherry* as described above and sequence-verified. Similarly, the *pENTR 1A- hobbit* entry vector was recombined into *pBID-UASC-G-mCherry* to generate *UAS-hobbit-mCherry* and was sequence-verified. To generate *UAS-hobbitΔN117-GFP, pENTR 1A-hobbit* was digested with *Kpn*I and *Age*I. The resulting ∼0.8 kb fragment was used as a template for PCR with the following primers: 5’-AAA GGT ACC CAA CAT GGC CTC GGA GGC GAA GGG TGT-3’ and 5’-5’AAA ACC GGT CGA TGT CCT CCG CTT GGC CT-3’. The resulting 0.37 kb fragment was digested with *Kpn*I and *Age*I and ligated into *pENTR 1A-hobbit*; the *pENTR 1A-hobbitΔN117* entry vector was recombined into *pBID-UASC-G-GFP* as described above and sequence-verified. All final plasmids were injected into *VK00027* flies for phiC31-mediated site-directed integration using standard techniques (Rainbow Transgenic Flies, Inc.). To generate *UAS-Stim*^*DDAA*^*-GFP* transgenic flies, plasmid LD45776 containing the cDNA sequence of *Drosophila Stim* Isoform A was obtained from the *Drosophila* Genomics Resource Center (DGRC). The *Stim* coding sequence was cloned by PCR using primers 5’-CAC CAT GCG AAA GAA TAC CAT TTG GAA C-3’ and 5’-TTC CGT GGC AAG CAG CGA AAA GTT C-3’ and ligated into *pENTR/D-TOPO* (Invitrogen). Site-directed mutagenesis (Stratagene QuikChange XL) was then used to convert the codons encoding D155 and D157 to codons for alanine using the primer 5’-GCT TGC ATC GTC AGC TAG CTG ATG CCG ATA ATG GAA ACA TCG-3’ and its reverse complement, followed by sequence confirmation. The *Stim*^*DDAA*^ coding sequence was then recombined into the pPWG destination vector (Carnegie *Drosophila* Gateway Vector Collection), which introduces a *UASp* promter and C-terminal EGFP tag, using LR Clonase. The final plasmid was injected into *w*^*1118*^ embryos for generation of transgenic animals by random transposable element insertion.

### Protein expression, purification, and lipid blots

The Apt1 fragment of the Hobbit protein (amino acids 1750-2300) and Apt1ΔC82 (amino acids 1750-2218) were amplified from the *pENTR 1A-hobbit plasmid* (described above) using the following primers: Apt1 F 5’- ATC CGG TAC CCA ACA TGG TAG TCT CAG AGA CTG TTG GAG CTT TCT TGA GCG AC-3’ and Apt1 R 5’-GCA TAA GCT TCT AAT GAT GAT GAT GAT GAT GGC TGC CGC CGC CGC CCA GCT CCT CCT TGC TCT CCC GG-3’; Apt1ΔC82 F 5’-ATC CGG TAC CCA ACA TGG TAG TCT CAG AGA CTG TTG GAG CTT TCT TGA GCG AC-3’ and Apt1ΔC82 R 5’-GCA TAA GCT TCT AAT GAT GAT GAT GAT GAT GGC TGC CGC CGC CGC CCC ACG TCA CAT TGT GAT ACT CCA GCG-3’. The reverse primers include a C-terminal five amino acid spacer and a 6x-His tag. The resulting PCR products were digested with *Kpn*I and *Hind*III, ligated into *pTRC99a* (Amann et al., 1988), and sequence-verified. Plasmids were transformed into *E. coli* EXPRESS BL21(DE3) chemically competent cells (Lucigen). A dense overnight culture was diluted 1:100 into 15 mL LB (Miller) with 100 µg/mL ampicillin. Cultures were grown at 37°C to A_600_ ∼0.4, then 200uM IPTG (final concentration) was added and cultures continued shaking for 3 h at 37°C. Cultures were then centrifuged for 5 min at 5000x*g*, and pellets were frozen at -80°C. For protein purification, pellets were thawed on ice for 15 min, resuspended in 3 mL lysis buffer (50 mM NaH_2_PO_4_, 300 mM NaCl, 10 mM imidazole, pH 8) with 1 mg/mL lysozyme (final concentration), 1.5 µL Benozoase nuclease (Sigma), and 1 µL protease inhibitor cocktail (Sigma P8849), incubated on ice for 30 min, and centrifuged for 30 min at 10,000x*g* at 4°C. 1.5 mL of Ni-NTA agarose (Qiagen) was washed three times with lysis buffer. Cleared lysate and washed Ni-NTA agarose were combined and incubated for 1 h at 4°C while rotating on a nutator. The slurry was loaded onto a gravity column, washed two times with 1 mL wash buffer (50 mM NaH_2_PO_4_, 300 mM NaCl, 20 mM imidazole, pH 8), and eluted four times with 0.5 mL elution buffer (50 mM NaH_2_PO_4_, 300 mM NaCl, 250 mM imidazole, pH 8). Proteins were visualized on 4-20% Mini-PROTEAN TGX precast gels (BioRad) and concentration was determined using the Bradford Protein Assay (BioRad). For lipid blotting experiments, Membrane Lipid Strips (Echelon Biosciences) were blocked in PBS with 0.1% Tween-20 (PBST)/3% BSA overnight while rotating at 4°C. The strips were then incubated for 1 h at room temperature with 1.5 mg of purified protein diluted in PBST/3% BSA to a 15 mL final volume. A 1:10,000 dilution of mouse α-(6X)His (Invitrogen) primary antibody in PBST/3% BSA was applied for 1 h at room temperature, followed by a 1:20,000 dilution of anti-mouse IgG HRP conjugated secondary antibody (Promega) for 1 h at room temperature. Washes with PBST were performed between each step. Lipid strips were then incubated for 5 min with SuperSignal West Pico PLUS Chemiluminescent Reagent (Thermo) and visualized using a UVP ChemiDoc-It^2^. Image brightness and contrast were optimized post-acquisition using Adobe Photoshop CS6.

### Confocal microscopy in Drosophila salivary glands

All images were obtained from live, unfixed tissues. Salivary glands were dissected from animals of the appropriate developmental stage and genotype in PBS and mounted in 1% low-melt agarose (Apex Chemicals) made in PBS. Tissues were imaged for no more than 15 min after mounting, and at least 10 salivary glands were imaged per experiment. Imaging was carried out at room temperature. Images were acquired using an Olympus FV3000 laser scanning confocal microscope (20x objective, NA 0.75 or 100x oil immersion objective, NA 1.49) with FV31S-SW software. The pinhole was closed down to 0.8 Airy disc for images acquired at high resolution (Fig. 4A, B, Fig. S3B). Images obtained as z-stacks, as indicated in the figure legends, were deconvolved using three iterations of the Olympus CellSens Deconvolution for Laser Scanning Confocal Advanced Maximum Likelihood algorithm. Brightness and contrast were optimized post-acquisition using Olympus FV31S-SW software. For imaging-based protease protection assays, salivary glands were adhered to a plastic coverslip in a minimal quantity of PBS; after at least 30 s of time-lapse image acquisition using the Olympus FV3000 resonant scan head, 50 µL of 0.05% digitonin (Invitrogen) was added, followed by an additional at least 1 min of imaging. Then we added 75 µL of 50 µg/mL proteinase K (Fisher Scientific), followed by at least 2 min of imaging.

## ACKNOWLEDGEMENTS

Reagents obtained from the Bloomington *Drosophila* Stock Center (NIH P40OD018537) and *Drosophila* Genomics Resource Center (NIH 2P40OD010949) were used in this study.

## AUTHOR CONTRIBUTIONS

Conceptualization: S.D.N., J.R.J., S.D.E., A.B; Methodology: S.D.N., J.R.J., S.D.E., A.B.; Validation: S.D.N., J.R.J., S.D.E., A.B.; Investigation: S.D.N., J.R.J., A.T.C., J.T.S., J.E.S.; Data curation: S.D.N., J.R.J.; Writing-original draft: S.D.N.; Writing-review and editing: S.D.N., J.R.J., J.T.S., S.D.E., A.B.; Visualization: S.D.N., J.R.J., S.D.E., A.B.; Supervision: S.D.E., A.B.; Project administration: S.D.E., A.B.; Funding acquisition: J.T.S., S.D.E., A.B.

---

**Figure S1.**
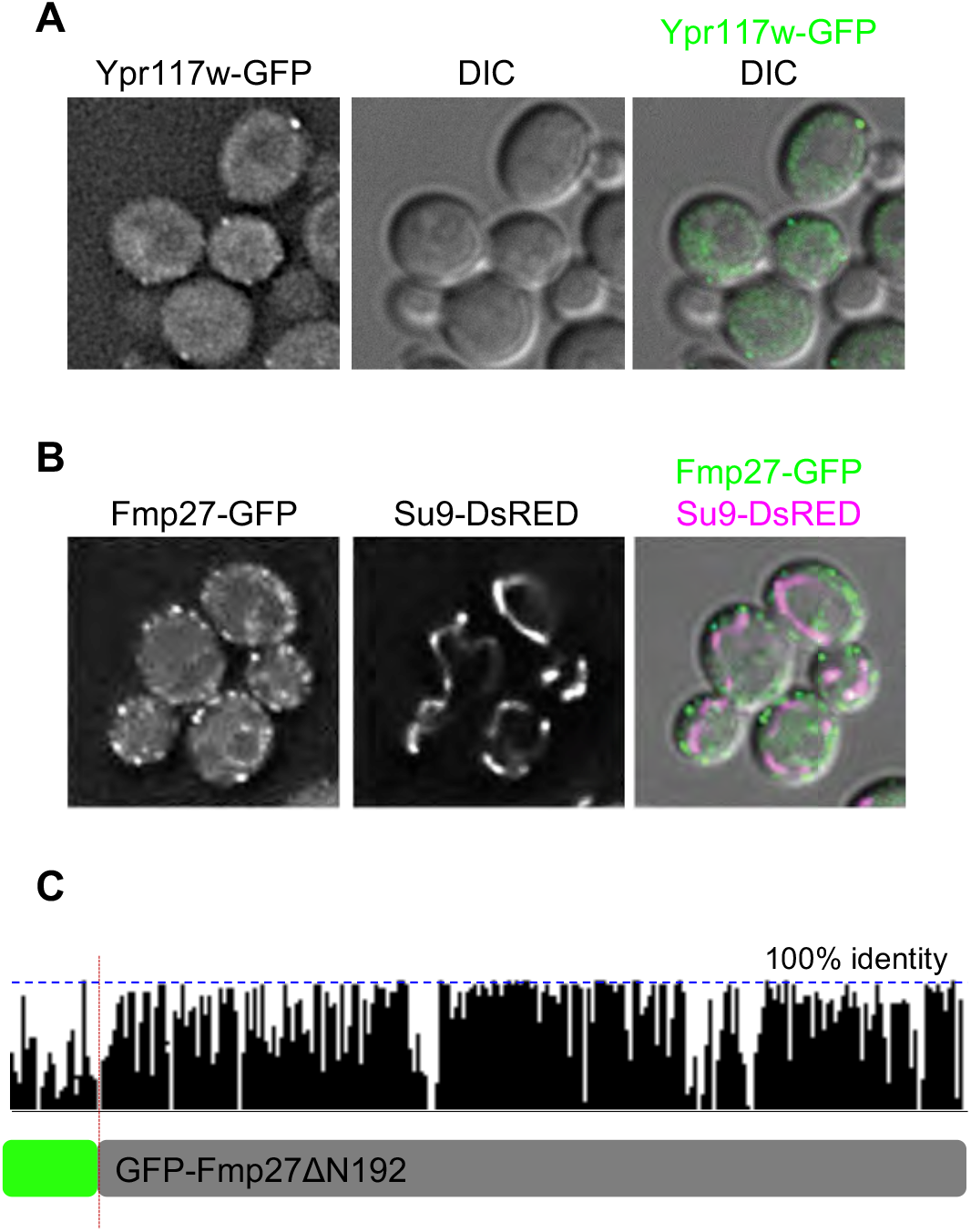
Analysis of Ypr117w and Fmp27 localization and conservation. **(A)** Live-cell imaging of endogenously tagged Ypr117w-GFP (green) shows that Ypr117w is enriched in puncta at the cell cortex; however, this protein was expressed at very low levels. **(B)** Live-cell imaging of endogenously tagged Fmp27-GFP (green) with the mitochondrial marker Su9-DsRED (magenta) shows that Fmp27 does not localize to mitochondria. Su9-DsRED was expressed from a plasmid. **(C)** Schematic depicting the conservation of *S. cerevisiae* Fmp27 and its orthologs in nine other fungal species (see methods section for list). GFP-Fmp27ΔN192 removes the first region of highly conserved sequence (192 amino acids) at the N-terminus.

**Figure S2.**
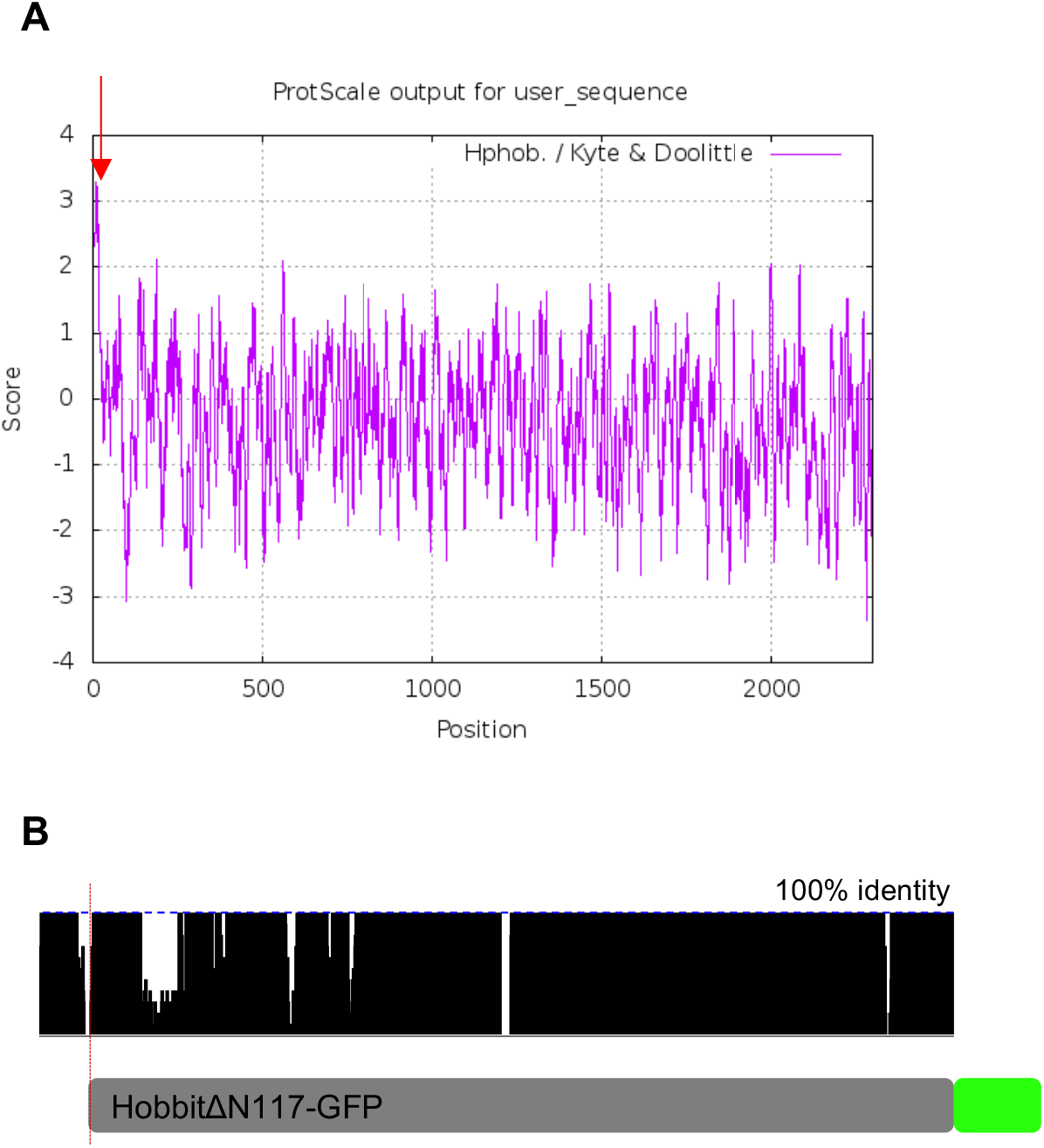
Design of *D. melanogaster* N-terminal Hobbit truncation. **(A)** Kyte-Doolittle hydrophobicity plot of *D. melanogaster* Hobbit shows a short, highly hydrophobic region at the N-terminus of the protein (marked by red arrow), consistent with a possible transmembrane domain. Plot generated using Expasy ProtScale. **(B)** Schematic depicting the conservation of *D. melanogaster* Hobbit and its orthologs in nine other Drosophilid species (see methods section for list). The HobbitΔN117-GFP transgene removes the first region of highly conserved sequence (117 amino acids) at the N-terminus.

**Figure S3.**
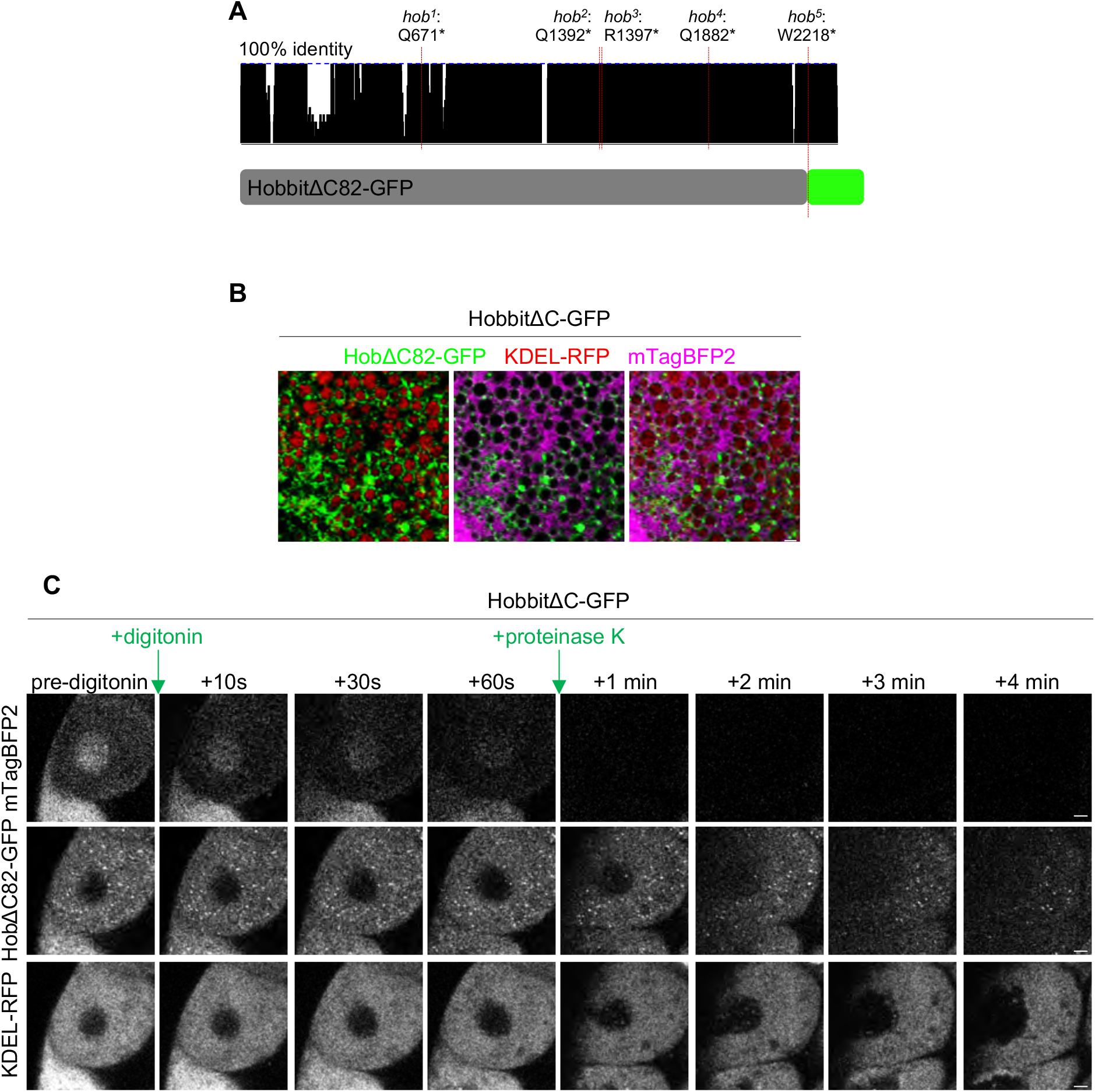
C-terminal truncation of Hobbit does not affect ER membrane localization or topology. **(A)** Schematic depicting the conservation of *D. melanogaster* Hobbit and its orthologs in nine other Drosophilid species (see methods section for list) and the position of each of the identified *hobbit* mutant nonsense mutations. The HobbitΔC82-GFP transgene enables overexpression of a protein comparable to that in *hob*^*5*^. **(B)** Live-cell imaging of HobbitΔC82-GFP (green), the ER lumen marker KDEL-RFP (red), and cytosolic mTagBFP2 (magenta) in salivary glands at the onset of metamorphosis (0 h after puparium formation) shows that C-terminally truncated Hobbit still localizes to the ER membrane. Full genotype: *UAS-KDEL-RFP/+; Sgs3>hobΔC82-GFP/UAS-mTagBFP2*. Images show a single slice from a z-stack comprising three optical sections at a 0.28 µm step size. **(C)** Imaging-based protease protection assay shows that the C-terminal truncation of Hobbit does not affect topology. Cytosolic mTagBFP2 (top) rapidly diffuses out of the cells after permeabilization with digitonin, while HobbitΔC82-GFP (middle) and KDEL-RFP (bottom) are unaffected. Like the full-length protein, HobbitΔC82-GFP (tagged at the C-terminus) is degraded after subsequent addition of proteinase K, while KDEL-RFP is unaffected. Note that the cells flatten after addition of proteinase K. Experiment was conducted using 0 h PF glands. Full genotype: *UAS-KDEL-RFP/+; Sgs3>hobΔC82-GFP/UAS-mTagBFP2*. Scale bar in (B): 1 µm; (C): 10 µm.

**Figure S4.**
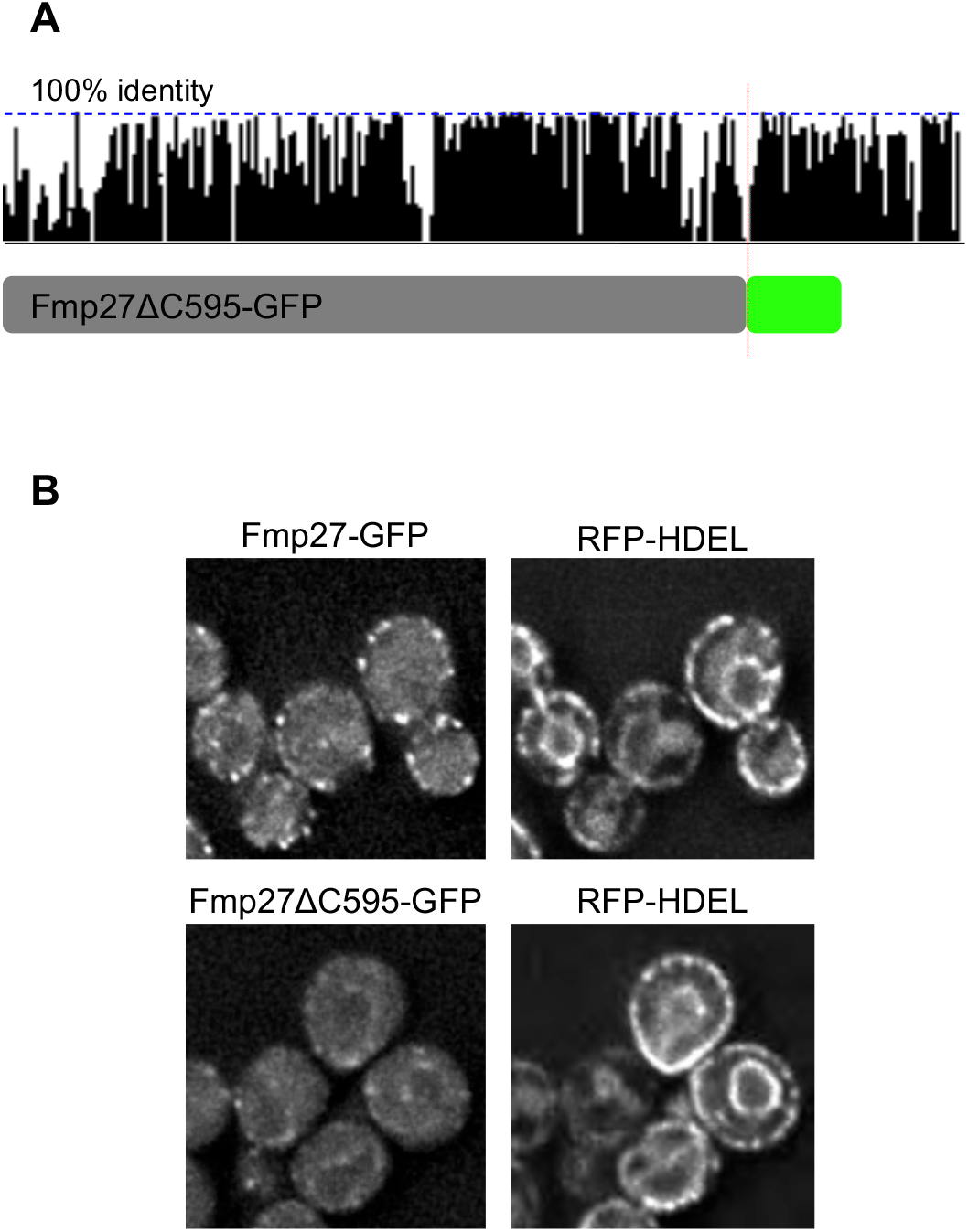
C-terminal truncation of Fmp27 reduces cortical ER localization. **(A)** Schematic depicting the conservation of *S. cerevisiae* Fmp27 and its orthologs in nine other fungal species (see methods section for list; also displayed in Fig. S1C). Fmp27ΔC595-GFP removes highly conserved sequences at the C-terminus (595 amino acids) **(B)** Live-cell imaging of full-length Fmp27-GFP and Fmp27ΔC595-GFP with the ER marker RFP-HDEL shows that C-terminal truncation of Fmp27 reduces its localization to cortical ER. FMP27- GFP and C-terminal truncation were made at the endogenous *FMP27* locus and are expressed under the endogenous promoter; RFP-HDEL was expressed from a plasmid.

**Movie 1. Imaging-based protease protection assay with full-length Hobbit-GFP**. Live-cell time-lapse imaging of cytosolic mTagBFP2 (cyan), full-length Hobbit-GFP (green), and KDEL-RFP (red) in an imaging-based protease protection assay. The selective detergent digitonin, which permeabilizes the plasma membrane but not intracellular membranes, was added at timestamp 28.049430 s, and proteinase K was added at timestamp 2 min 4.664132 s. Still images from this time-lapse are pictured in Fig. 4C.

**Movie 2. Imaging-based protease protection assay with HobbitΔC82-GFP**. Live-cell time-lapse imaging of cytosolic mTagBFP2 (cyan), HobbitΔC82-GFP (green), and KDEL-RFP (red) in an imaging-based protease protection assay. The selective detergent digitonin, which permeabilizes the plasma membrane but not intracellular membranes, was added at timestamp 1 min 45.964512 s, and proteinase K was added at timestamp 3 min 47.512041 s. Still images from this time-lapse are pictured in Fig. S3C.

